# Robotics-driven manufacturing of cartilaginous microtissues for the bio-assembly of skeletal implants

**DOI:** 10.1101/2023.01.09.522841

**Authors:** Isaak Decoene, Gabriele Nasello, Rodrigo Furtado Madeiro de Costa, Gabriella Nilsson Hall, Angela Pastore, Inge Van Hoven, Samuel Ribeiro Viseu, Catherine Verfaillie, Liesbet Geris, Frank P. Luyten, Ioannis Papantoniou

## Abstract

Automated technologies are attractive for enhancing a robust manufacturing of tissue engineered products for clinical translation. In this work, we present an automation strategy using a robotics platform for media changes of cartilaginous microtissues cultured in static microwell platforms. We use an automated image analysis pipeline to extract microtissue displacements and morphological features, which serve as input for statistical factor analysis. To minimize microtissue displacement and suspension leading to uncontrolled fusion, we performed a mixed factorial DoE on liquid handling parameters for large and small microwell platforms.

As a result, 144 images, with 51 471 spheroids could be processed automatically. The automated imaging workflow takes 2 minutes per image, and it can be implemented for on-line monitoring of microtissues, thus allowing informed decision making during manufacturing. We found that time in culture is the main factor for microtissue displacements, explaining 10 % of the displacements. Aspiration and dispension speed were not significant at manual speeds or beyond, with an effect size of 1 %. We defined optimal needle placement and depth for automated media changes and we suggest that robotic plate handling could improve the yield and homogeneity in size of microtissue cultures. After three weeks culture, increased expression of COL2A1 confirmed chondrogenic differentiation and RUNX2 shows no osteogenic specification. Histological analysis showed the secretion of cartilaginous extracellular matrix. Furthermore, microtissue-based implants were capable of forming mineralized tissues and bone after four weeks of ectopic implantation in nude mice.

We demonstrate the development of an integrated bioprocess for culturing and manipulation of cartilaginous microtissues. We anticipate the progressive substitution of manual operations with automated solutions for manufacturing of microtissue-based living implants.

## Introduction

Spheroids are small 3D structures built by spontaneous aggregation of single cells and they are increasingly used both for drug screening applications (1) and in the field of regenerative medicine (2). As the cells produce their own extracellular matrix and microenvironment, the spheroids can be considered microtissues (3). When complex organisation and function appears, the term organoid can be used (4,5). Compared to 2D monolayer cultures, 3D cultures allow cell growth, matrix deposition, and matrix organisation in all directions, which is a more natural environment for cells. In tissue engineering applications, they are ideal as building blocks to create large and complex tissues from the bottom up (6). Because of their small size, there are no diffusion limitations on nutrients or growth factors, allowing a more precise control of differentiation (7). The current bottleneck in this field is the creation of tissues with sufficient volume and a robust quality profile. To further harness the properties of these cellular building blocks, scalable manufacturing and the production in a controlled manner is required to ensure a predefined quality profile.

Microtissue production is achieved by means of different methods including the hanging drop method, drop-seeding cells on low adherence substrates, spinning bioreactors, microfluidics, magnetic aggregation, or the use of non-adherent microwells as reviewed in Liu et al. (4). Dynamic cultures are easily scalable, but harsh on the cells and microtissue size control is difficult (8,9). The use of biomaterials both for the creation of spheroids and for downstream steps allows versatile and controlled tissue production, differentiation, and growth (10). Hydrogels can be produced in-house with tuneable shapes, sizes, mechanical and biological properties, making it an attractive option for research purposes (11). Due to a limited number of relevant clinical grade biomaterials, the use of sacrificial hydrogels (12) or scaffold-free approaches are also promising. However, these platforms lack scalability, and are prone to production errors.

On the other hand, many commercial microwell systems are available, including EZSPHERE™, AggreWell™, Elplasia plate, SpheroFilm, SphericalPlate 5D, and 3D Petri Dish^®^. Currently, microtissue production in microwell platforms requires manual handling and pipetting, which is prone to errors, including microtissue escape from microwells, followed by uncontrolled agglomeration and thus, batch failure. Monitoring of microtissue cultures and their morphometric quality profiles can be done non-invasively through imaging. Software is already available to segment and analyse microtissues in hydrogel microwells and floating cultures, even enabling selection of desirable microtissues and studying their fusion kinetics (13–17). Commercial microwell systems are typically produced from PCL or PDMS in different microwell shapes, which interact with light, creating complex backgrounds in brightfield images. Manual segmentation using for example fiji (18) or napari (19) is time consuming and person-dependent, but well suited for small datasets and varying input images (20–23). Automated segmentation is fast, reproducible and can be done with minimal supervision. Yet, for complex images we still need methods for automated segmentation to monitor microtissue cultures.

Recently, the use of chondrogenic microtissues and organoid assemblies has shown promising results in long-bone defect regeneration through endochondral ossification following the paradigm of developmental engineering (24–26). Periosteum derived cell aggregates form transient cartilage microtissues in chondrogenic medium (27). Bone forming potency was shown ectopically in nude mice for both individual spheroids as larger constructs and a proof-of-concept was provided for successful healing of a murine critical-sized long bone defect (28). The critical next step is a bridging the gap towards large animal models in a preclinical phase. For long bone defects, ovine models are well suited regarding biological similarity, long bone dimensions and mechanical loading during normal behaviour (29). A transition to preclinical studies and industrial translation requires a significant scale-up of this bottom-up strategy beyond what is possible by manual methods.

A well-characterised process and manufacturing line is imperative towards a successful translation (Rousseau et al., 2018, ten Ham et al., 2018). Significant progress has been made in automated cell culture, passaging and organoid seeding (31–34), showing the feasibility and advantages of automating manual processes. Furthermore, robotic platforms using microtissues to create complex tissue constructs are appearing (35,36). Yet, robotic production of microtissues for these applications is underexplored. In this study, we explored the integration of robotics to automate media changes and imaging for cartilaginous microtissue differentiation in parallel with automated analytics essential for robust manufacturing of TE ATMPs (37–39). Designs of experiment (DoE), as goal-oriented statistical approaches to defining factor importance towards predefined critical quality attributes have been used successfully in TE research (27,40,41). In this work, we combine statistical DoE approaches with automated image segmentation and analytics to optimise robotic media changes of cartilaginous bone-forming spheroids in static microwell culture platforms.

## Results

### Automated image analysis pipeline

We created a pipeline connecting machine learning-based image segmentation software (ilastik(42)) and object detection and data extraction software (cellprofiler(43)) to statistical data analysis scripts (R). These different modules are connected through jupyter notebooks written in python. All software is freely available and is gathered in one Docker container.

The process, described in **figure 2a**, starts with images containing any number of microtissues in square-shaped microwells. The images must be taken with identical settings to generate optimal results. In a first step, a pixel classification algorithm is trained in ilastik to predict whether a pixel belongs to a microtissue or background, resulting in a probability map. Then, a second pixel classification algorithm is trained to generate a mask of the microwell pattern. In cellprofiler, the microtissue probabilities are thresholded to identify separate objects. These objects are then described in 140 morphometric parameters including object shape (e.g., location, area, diameter, roundness) and texture (e.g., intensity variation, pixel entropy and correlations). Individual microwell contours are identified from the microwell pattern mask. For each microtissue, we calculated whether it is in a microwell, as a child-parent relation. As a result, we could calculate which microwells are empty, and which contain more than one microtissue. To finish, a virtual image is plotted containing the microtissues respective location and size. Moreover, as shown in **supplemental figure 2**, a density plot shows the size distribution within one image. Through a jupyter notebook, this pipeline can be used in batch mode to process hundreds of images containing thousands of microtissues. The data presented here contains information from 144 individual images, and 51 471 microtissues. Training the machine learning algorithm for pixel classifications required a desktop with 64GB RAM and 16 cores, while images were classified in a laptop with 8GB RAM and 2 cores. The computing time to classify, segment and extract quantitative parameters from new images was 2 minutes per image. Assuming a spherical shape in three dimensions, we can estimate microtissue volume from the two-dimensional projection. With this approach, we performed grain analysis for the size distribution of individual microtissues over time as shown in **table 1**. On average, we found that microtissues increased in size over time in both platforms. The variation in size, which is represented as the span measurement, indicates less size variability for the large microwells. Furthermore, the total volume produced in one well plate is calculated as the sum of microtissue volumes, corrected for the percentage of microtissue displacement. We found that large microwell platforms generate a larger tissue volume with less microtissues.

**Table 1:** Summary statistics on microtissue size distribution.

| Platform | Time | Microtissues measured | μwells | Not empty (%) | μtissue volume ($\text{mm}^3$) | D10 ($\text{mm}^3$) | D50 ($\text{mm}^3$) | D90 ($\text{mm}^3$) | Span | Predicted tissue volume ($\text{mm}^3$) |
| --- | --- | --- | --- | --- | --- | --- | --- | --- | --- | --- |
| Aggrewell400 | day 7 | 14867 | 28800 | 91.1809 | 0.0015 | 0.0009 | 0.0017 | 0.0033 | 1.4571 | 42.1512 |
|  | day 14 | 13210 | 28800 | 83.6241 | 0.0017 | 0.0010 | 0.0020 | 0.0054 | 2.2674 | 49.8173 |
|  | day 21 | 12647 | 28800 | 78.8870 | 0.0021 | 0.0012 | 0.0024 | 0.0070 | 2.4076 | 59.1178 |
| Aggrewell800 | day 7 | 4014 | 7200 | 89.3833 | 0.0059 | 0.0033 | 0.0068 | 0.0160 | 1.8612 | 42.5603 |
|  | day 14 | 3574 | 7200 | 73.2044 | 0.0066 | 0.0036 | 0.0084 | 0.0162 | 1.5071 | 47.4942 |
|  | day 21 | 3158 | 7200 | 70.7658 | 0.0093 | 0.0048 | 0.0118 | 0.0318 | 2.2735 | 66.9176 |

### Automation and optimization of liquid handling parameters

One of the greatest challenges in long term culture of microtissues in static well plates is their sensitivity to liquid handling. The microtissues can easily be displaced, leading to premature uncontrolled fusion into larger irregular tissues that complicate downstream processes. We used the automated liquid handling station as shown in **figure 1** to maximize dispension and aspiration speeds for both small (aggrewell400) and large (aggrewell800) microwell platforms. In this way, the automated liquid handling station minimized microtissue displacement and time needed for media changes.

**Figure 1.**
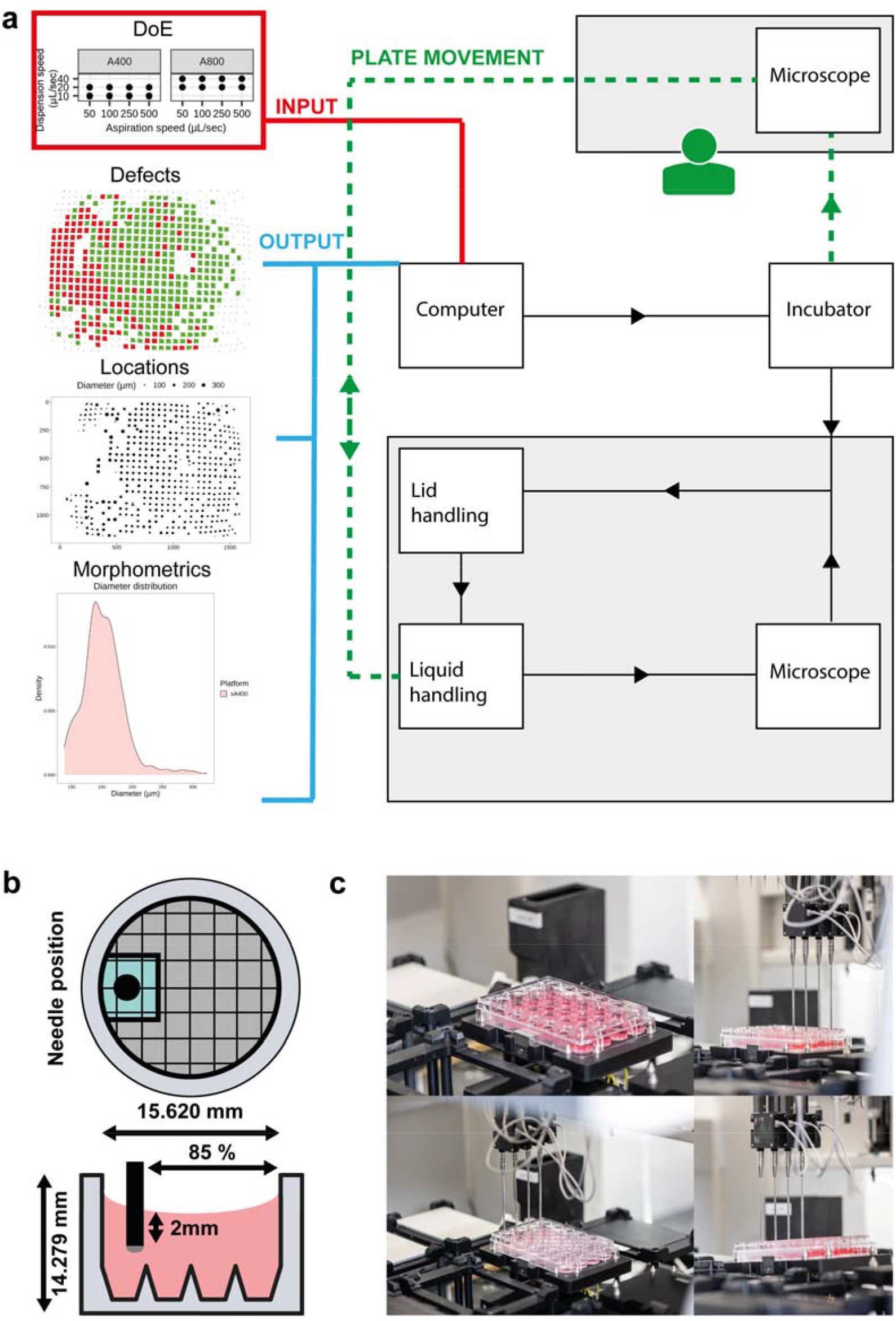
Setup for automated media changes. (a) Schematic pipeline representation starting from experimental designs as input leading to the automated generation of quality output. (b) Positioning of the liquid handling needle in a well plate. (c) Images of the liquid handling station during media changes of 24-well plates.

In a first design of experiment (DoE), we assessed the effect of aspiration speed, dispension speed and needle depth in aggrewell400 through a 2^3^ full factorial design as shown in **supplemental figure 1a-e**. In this experiment, we found no significant (p < 0.05) effects for the tested factor levels, but the data showed an indicative effect (p < 0.1) for both the dispension speed alone as the interaction of dispension speed with needle depth (supplemental **figure 1e**). Both aspiration speed and needle depth were not significant in the range tested, but we noticed dripping for a 1mm needle depth. Parameter testing requires a high amount of living microtissues, medium and time. However, we found that gelatine beads that are commonly used as a growth surface in suspension cultures, can be used as substitutes for living microtissues (**supplemental figure 1g**). Because of the observed interaction between dispension speed and needle depth, we further optimized needle position by varying the needle depth and the distance from the edge of the well (**supplemental figure 1f**). Because of the surface tension, the liquid has a concave shape. As a result, the needle can be place further from the microtissues when placed more towards the side of the well. However, at 90 % or more, the needle touches the well plate, disrupting the entire 24-well plate. An optimum was reached by placing the needle 2 mm below the liquid surface at 85 % from the well centre. As the needle follows the liquid level, a minimal number of spheroids were moved.

A second DoE with two mixed factorial designs was set up as shown in **figure 3a**. As we suspected the A800 to be less sensitive, we decreased the dispension speed of the A400, but kept the same for A800. Aspiration speed was not significant in the first DoE, so we increased the speed up to 500 μL/sec. On day 7, 14 and 21 brightfield images were taken to assess microtissue displacement and the cumulative effect over time. The automated image analysis pipeline was used for segmentation and data extraction as explained in **figure 2**. The main effects in **figure 3b-e** show a significant average increase in defects over time for both platforms. Dispension speed has no significant effect, but in A800, a significant effect was observed for aspiration speed. Especially for 100 μL/sec, a high defect percentage was measured. However, upon inspection of the defect distribution in the 24-well plate shown in **figure 2b**, most defects locate in the bottom half of the well plate. As seen from the interaction effects in **figure 3k**, the 100 μL/sec condition in the A800 platform is not in line with higher or lower aspiration speeds.

**Figure 2.**
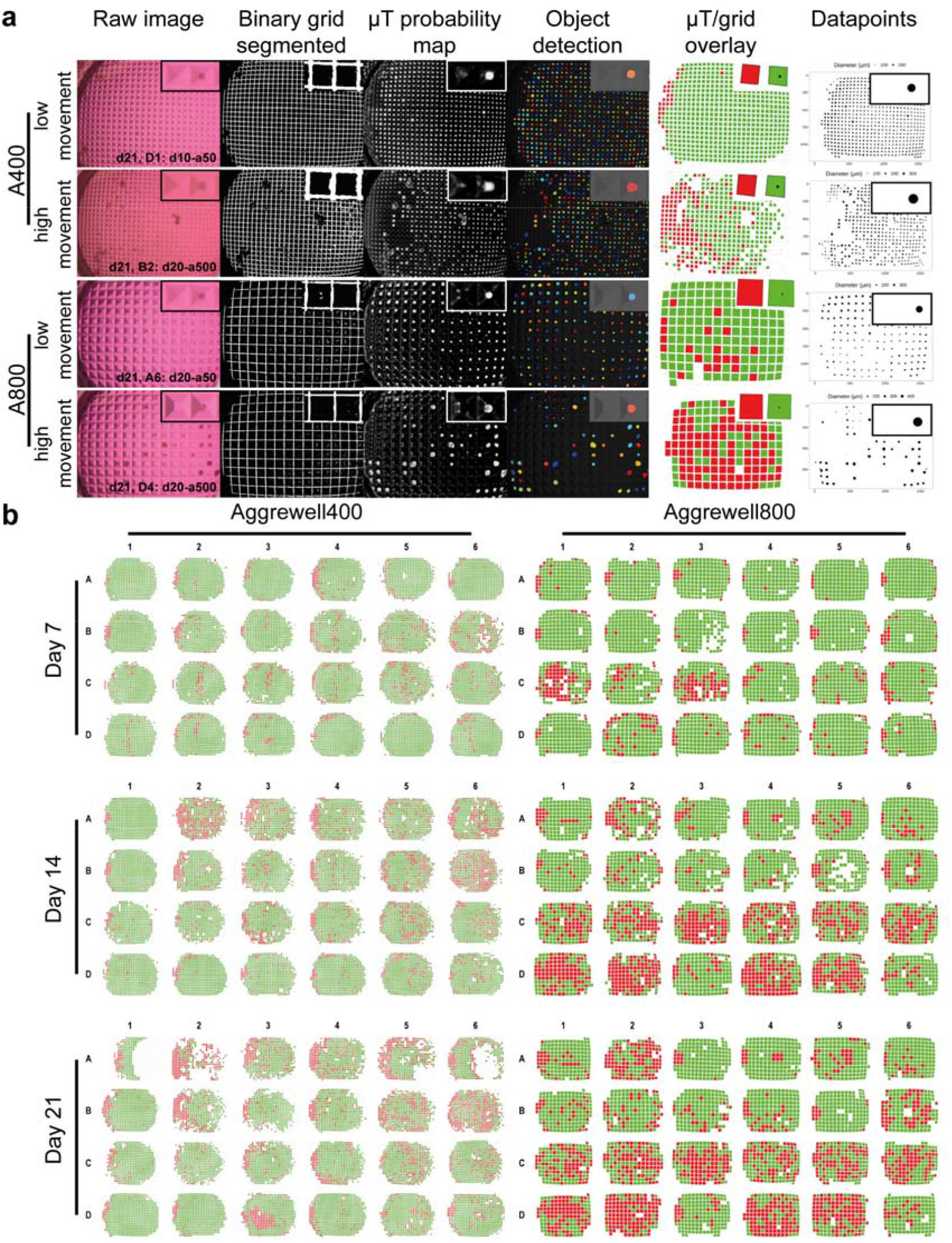
Image analysis pipeline. (a) Representative images of large and small microwells with high and low microtissue movement. For each image, the microwell grid was segmented and a microtissue probability map of all pixels was generated. Individual spheroid objects were identified and overlayed on the microwell grid. Each microwell is coloured red (containing no microtissue) or green (contains one or more microtissues). Finally, each microtissue is converted to a datapoint, here represented by its location and diameter (μm). (b) An overview of all 144 images that were analysed, showing microtissue displacements.

**Figure 3.**
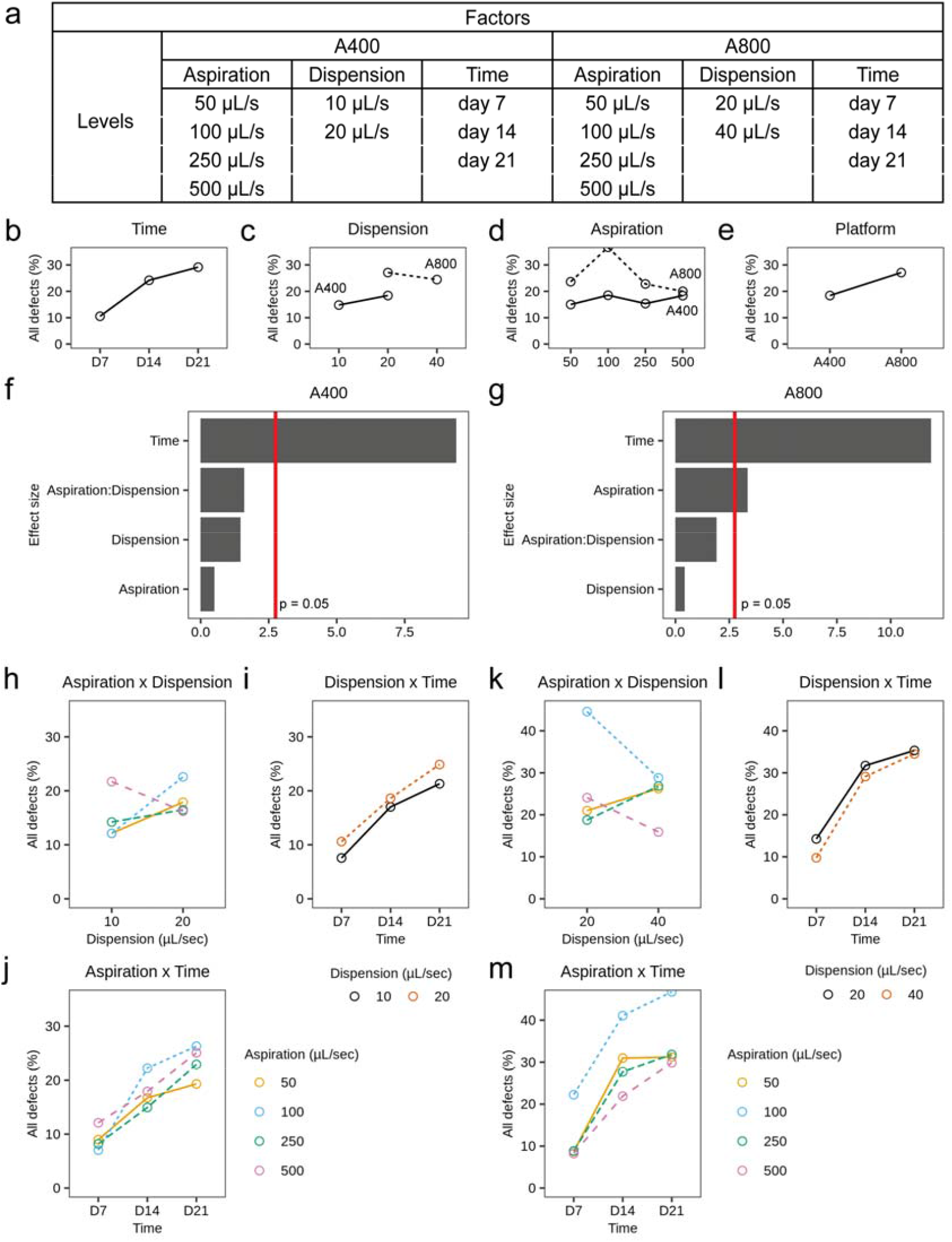
Design of Experiment (DoE) setup and results. (a) Experimental design with the tested factors levels. (b-e) Main effects plots. (f-g) Statistical significance of main and interaction effects for A400 and A800 platforms. (h-j) Interaction plots for A400, and (k-m) interaction plots for A800.

### Differentiation potential of sheep PDCs

To assess the differentiation potential of the sheep periosteum-derived cells, gene expression analysis and histological stainings were performed on day 21 and compared to day 0 monolayer culture in growth medium. **Figure 4a** recapitulates the timeline. In **figure 4e-f**, we see a downregulation of proliferation and progenitor genes, CD200 and PDGF. At the same time, **figure 4g-k** shows that chondrogenic markers BMP2, BMP4, TGFb1, and COL2A1 were upregulated while osteogenic marker RUNX2 was downregulated in both platforms. Only TGFb1 showed a significant difference between the aggrewell400 and aggrewell800 platform. Also, the DNA content shown in **figure 4l** after 21 days is the same for both platforms. We then analysed the microtissues on a tissue-level through (immune)histochemistry (**figure 4b-d)**. Cells are homogenously present throughout the microtissue with a thin layer on the outside. Alcian Blue staining shows the secretion of cartilaginous glycosaminoglycans in the extracellular matrix (**figure 4c**), but the degree of sulphation, shown in **figure** 4d, was higher in larger microtissues.

**Figure 4.**
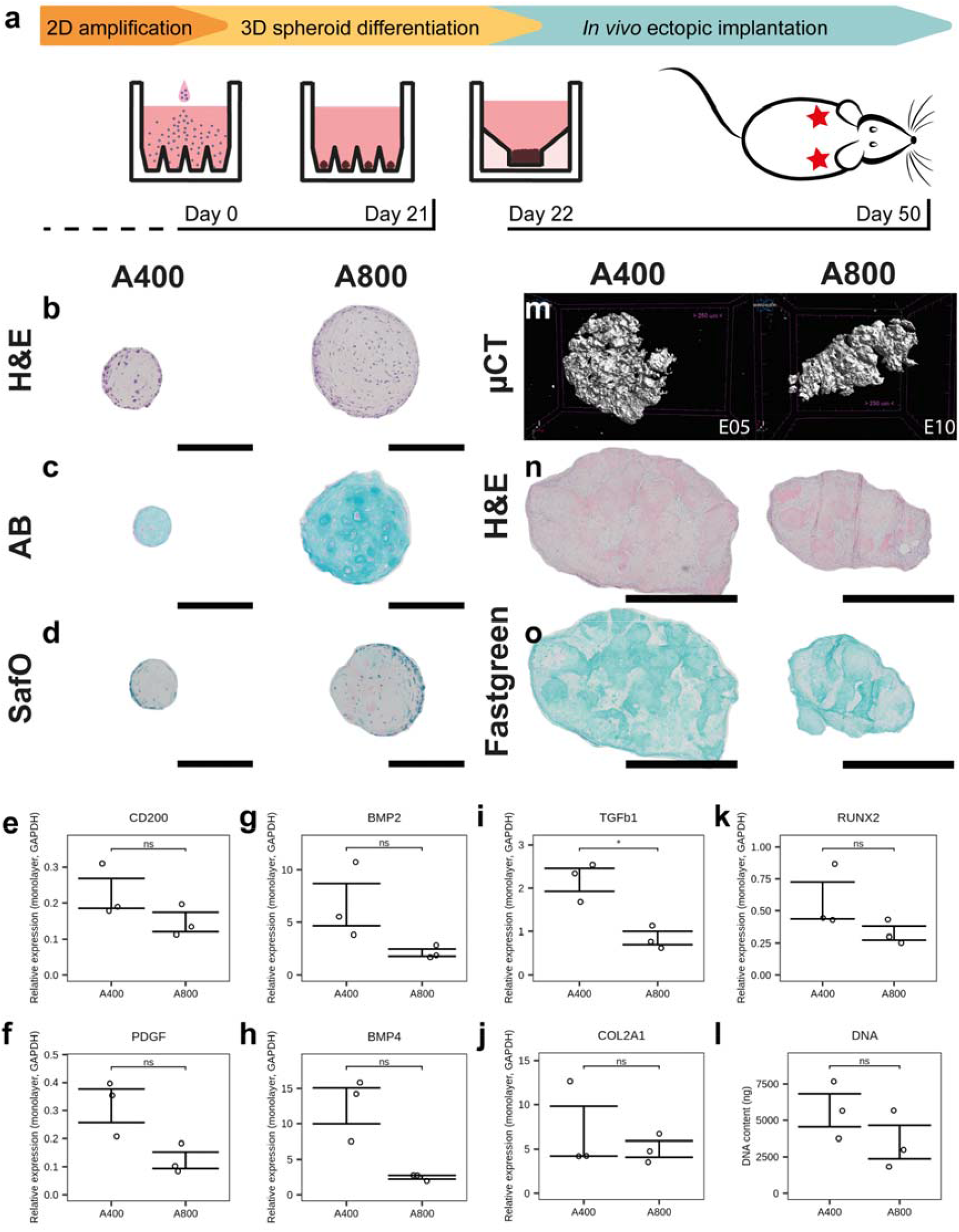
Biological characterisation of microtissues and ectopic explants. (a) Protocol overview and timeline. (b-d) histological staining of day 21 microtissues showing (b) Haematoxilin-Eosin, (c) Alcian Blue, and (d) Safranin O. Scale bars = 200 μm. (e-k) mRNA quantification of differentiation markers. (l) DNA quantification of individual wells on day 21. (m) Mineralisation after 4 weeks ectopic implantation of microtissue constructs measured by microCT analysis. (n) Haematoxilin-Eosin, and (o) SafraninO-Fastgreen staining of representative explants. Scale bars = 1 mm.

After 21 days, microtissues were fused in one construct and implanted ectopically for 4 weeks in nude mice to assess bone forming potential. Microcomputed tomography (μCT) shows mineralization of all implants, but no cortical bone formation or bone marrow compartments (**figure 4m**). H&E staining shows remainders of microtissue shapes that are not yet remodelled (**figure 4n**). Yet, the explants stain positive for Fast Green (**figure 4o**), consistent with the mineralization seen in μCT.

## Discussion

Using cellular aggregates, spheroids, microtissues and organoids as building blocks for the bottom-up engineering of progressively larger tissues is becoming a distinct domain in Tissue Engineering (44,45). While this strategy shows promise, the development of a robust biomanufacturing process is hampered by complex manual protocols, lack of scalability, and limited non-invasive quality measurements. To industrialise cell therapy and regenerative medicine, biomanufacturing of robust processes will lead to standardization and reproducibility while further enhancing process capability and cost efficiency (46–48). The implementation of Quality by Design principles requires technology developments towards measurable microtissue characteristics that are linked to the final product quality. Bottom-up engineering allows the early embedding of quality aspects already in the microtissues which are then incorporated in larger tissues. This is an important step for realising the potential for microtissue based TE Advanced Therapeutics (49).

In this work, we present an automation strategy for the differentiation of cartilaginous microtissues by combining robotic media changes with automated brightfield-image based analysis and multifactorial design of experiments for the optimization of this differentiation process. It has previously been shown that automating methods using robotics is an efficient way to reduce human error and operator-dependent variability in laboratory operations (50). One of the challenges for culturing microtissues in non-adherent well plates, is that over time they are prone to move, leading to uncontrolled fusion and the development of large agglomerates (23). This can result in tissue heterogeneity and influence the homogeneity of differentiation cascades over time leading also to the development of non-differentiated tissue locations (17) By introducing robotic medium changes, we investigated the effect of dispension and aspiration speed in ranges close to and beyond manual speed and found no statistically significant effect. Therefore, robotic media changes can be time saving. We did detect a significant time dependence, which we attribute to the manual handling of the microwell plates between incubator, microscope, and media change station. Fully robotic plate handling and imaging is thus considered to be the next step as we expect it could allow continuous production of microtissues.

To date the adoption of robotics for cell therapy and regenerative medicine applications has been carried out for the expansion of single cell populations such as adult MSC expanded in bioreactor systems (34). Moreover, robotic based expansion and differentiation to induced pluripotent stem (IPS) cells towards retinal pigment epithelial cells showed a high level of cell purity and functionality (51,52). However, we believe that in this work we tackle a bioprocess problem related to non-adherent bioprocesses and solutions aiming at the integration of robotic technologies in streamlined manufacturing pipelines need to be developed.

To enable full process control and automation, sensor technologies need to be integrated. Automated Imaging technologies have been implemented to characterize 2D cell culture processes such as expansion of MSCs (53,54) but also for the automated imaging of spheroid and microtissue structures (13–16,23,55) The use of automated image analysis contributed to further minimizing human error and variability which could also greatly improve the manufacturing process as it feeds back to the process parameters used during manufacturing (56). For instance, larger microwells produce less microtissues per area, but the total volume produced is higher. As these microtissue populations serve as active raw materials for living implants, morphological properties are important considerations for bio-assembly methods (57,58).

The automated classification of microtissues was achieved with a supervised machine learning algorithm. Compared to deep learning methods, ilastik’s random foster classifier required a significantly smaller training dataset (42), which was sufficient to reach correct classification. For this specific application, training the classifier required a desktop with higher computing power than a regular office laptop, however, the classification of new brightfield images was is computationally demanding. The entire automated imaging pipeline takes less than 2 minutes, and it has been isolated in a Docker container (59). Thus, the imaging workflow can be implemented in the same laptop used to acquire the brightfield images, allowing for live monitoring of the microtissues.

In addition, the integration of process analytical technologies (PAT) (60–63) and the definition of biological critical quality attributes by morphometrics, spectroscopic techniques or secreted proteins could be a next step to ensure a predictive outcome upon implantation (64–69). On line, non-invasive monitoring techniques, such as the current approach, can produce large quantities of data on the process and biology that can be used as input for machine learning algorithms. For example, users can train a classifier to identify patterns in the segmented microtissue images for predictive maintenance (70). A machine learning classifier can predict the failure of the microtissue manufacturing process and support informed decision making in the early stages of cell culture. Ultimately, this can help process optimisation to achieve full process control (71–74).

Finally, regarding the biological results obtained in this work, we used periosteal cells from sheep since they are frequently used as large animal models for preclinical studies targeting the healing of long-bone fractures and would be the animals of choice to validate bone tissue engineered products before the transition to a clinical trial (75). We see encouraging differentiation and the formation of cartilaginous microtissues that mineralize upon implantation showing regions of bone with blood vessel invasion and small areas of bone marrow. Further investigation of these implants at orthotopic sites, such as a critical size segmental tibial defects, will be required in future studies.

In conclusion, a stepwise automation of manual tasks, and eventual fully automated biomanufacturing of cartilaginous microtissue populations is becoming increasingly feasible, enabling next steps towards the translation process. Here, we show how robotic media changes combined with statistical DOE, non-invasive imaging and automated image analysis can be used to optimize large-scale biomanufacturing of microtissue building blocks, while being compatible with non-invasive quality monitoring.

## Material and methods

### Cell expansion

Periosteum biopsies were obtained from sheep tibia. After digestion, the cells were cultivated for 8 passages in expansion medium containing DMEM (gibco) supplemented with 1 % antibiotic– antimycotic (Invitrogen) and 10 % FBS (South Afrika FBS, BioWest, France).

### Microtissue formation

The commercially available microwell platform (AggreWell™800 or AggreWell™400, STEMCELL Technologies Inc, Canada) was coated with Anti-Adherence Rinsing Solution (STEMCELL Technologies Inc, Canada) to avoid cell attachment, centrifuged to ensure homogenous coating and washed with basal medium prior to cell seeding. Sheep periosteum derived cells (sPDCs) were harvested with TrypLE Express (Life Technologies, UK) and seeded at 300 000 cells per 2 mL chondrogenic medium resulting in a microtissue size of 250 or 1000 cells for aggrewell400 and aggrewell800 respectively. The cells self-aggregate and were differentiated for 21 days in a serum-free chondrogenic medium (CM) containing low glucose DMEM (Gibco) supplemented with 1% antibiotic– antimycotic (Invitrogen), 1 × 10^−3^ M ascorbate-2 phosphate, 1 × 10^−7^ M dexamethasone, 40 μg mL^−1^ L-proline, 20 × 10^−7^ M of Rho-kinase inhibitor Y27632 (Axon Medchem), ITS+ Premix Universal Culture Supplement (containing 6.25 μg mL^−1^ insulin, 6.25 μg mL^−1^ transferrin and 6.25 ng mL^−1^ selenious acid, 1.25 μg mL^−1^ bovine serum albumin (BSA) and 5.35 μg mL^−1^ linoleic acid, Corning), 100 ng mL^−1^ BMP2 (INDUCTOS), 100 ng mL^−1^ GDF5 (PeproTech), 10 ng mL^−1^ TGF-β1 (PeproTech), 1 ng mL^−1^ BMP-6 (PeproTech), and 0.2 ng mL^−1^ basic FGF-2 (R&D systems). Half of the medium was changed on day 3, 7, 10, 14 and 17.

### Robotic handling, automated medium changes and imaging

The STEm celL Laboratory Automation (STELLA) platform, funded by the NextGenQBio Hercules Foundation grant, was used to create and perform automated protocols for medium change and imaging. It is equipped with two liquid handlers (Biomek NX MC and Biomek NXp – Span8) (Beckman Coulter), one SCARA robotic arm, one Cytomat 10C Automated Incubator and one Cytomat Microplate Hotel (Beckman Coulter), one High-content Imaging System Image eXpress (Molecular Devices), two Plate Delliders (Beckman Coulter), one CapitAll IS Automated Capper/Decapper (Thermo Fisher), one Asymptote Freezer (Grant) and one Sigma 6K15 Centrifuge (Sigma), all inside a BSL2 sterile enclosure. This system enables our group to perform fully automated plate handling, medium change and imaging based on a design of experiment (DoE) to explore the best conditions for automated pipetting in which the microaggregates are not displaced or aspirated. Briefly, Aggrewell plates containing the microaggregates were manually placed in the incubator 24 hours prior the automated sequence starts. Plates were then manually moved from the incubator to the Span8 liquid handler where different liquid aspiration and dispensing, needle height and position were tested according to the DoE. In between the first and second DoE, needle placement was optimized using gelatin microcarriers (CultiSpher S, Percell), which have similar characteristics as living microtissues.

### Automated image analysis

Brightfield images of microtissues cultured in Aggrewell™ were taken manually with an inverted DMi1 microscope (2.5x, 0.07 NA lens, Leica). A custom-made image processing workflow was implemented to automatically segment microtissues and extract quantitative morphological information.

Microtissue segmentation was performed by the pixel classification algorithm of ilastik(42) software (v. 1.3.3). Brightfield images were annotated with two classes (microtissues and background). 1 image per timepoint per condition was annotated to train the default ilastik classifier (Random Forest with 100 trees and all image filters selected). The training dataset for pixel classification consisted of 4 images where a total of 80 microtissues were manually labelled according to a specific pattern. Images were divided in 4 quadrants and the pixels were assigned to the two classes for 6 wells of each quadrant.

The ilastik classifier generated probability maps of each class that were converted to .tif files and transformed into individual objects in cellProfiler(76) software (v 4.2.1). First, individual (primary) objects were identified by global Otsu thresholding of the microtissue class and declumping to distinguish touching objects. Later, the identified objects were filtered by diameter, compactness, eccentricity and area to remove small or irregular objects (76). Finally, several morphological features of the filtered objects (i.e., area and shape) were measured.

Besides segmenting microtissues, another image processing workflow was run in parallel to detect the microwells and localize microtissues in the culture plate. The ilastik classifier was also trained to segment the microwell boundaries from the background. The probability maps generated by ilastik were analyzed with the Python library OpenCV (77) detect the boundary points of each microwell. Then, the sp package in R(78) was used to match microtissue locations to microwell locations. Microwells were considered empty if the centroid of no microtissue laid within the boundary points.

### Formation of macro-construct

Custom round-bottom macrowells were created in 3 % agarose (w/v) (Invitrogen) and sterilized using UV. Microtissues from 3 wells were gently flushed out from their microwells on day 21, concentrated and added to the macrowells to sediment for 1 hour. Chondrogenic medium was added, and the constructs were incubated for 24 hours at 37 °C, 5 % CO2 and 95 % humidity to fuse into a coherent implant.

### *In vivo* ectopic implantation

An incision was made on the back of the mice under general anaesthesia (ketamine/xylazine) to create two pockets at the shoulder region per mouse. Implants were removed from the agarose macrowell, washed in 1xDPBS to remove loosely bound growth factors. 4 constructs per condition were implanted subcutaneously in the back at the shoulder region of 4 different female immune compromised mice (Rj:NMRInu/nu), with one implant per condition per mouse. The incision was closed by staples followed by postoperative administration of buprenorphine as pain relieve. After 4 weeks, the mice were sacrificed by cervical dislocation. Afterwards, implants were taken out and fixed for 4 hours in 4 % PFA.

### Nano-CT

3D quantification of mineralised tissue in PFA-fixed explants was done through nano-CT (Pheonix Nanotom M, GE Measurement, and Control Solutions). Explants were scanned with a diamond target, mode 0, 500 ms exposure time, 1 frame average, 0 image skip, 2400 images, and a 0.1 mm aluminium filter. Samples were scanned with a voxel size of 3 μm. CTAn (Bruker micro-CT, BE) was used for all image processing and quantification of mineralized tissue based on automatic Otsu segmentation, 3D space closing, and despeckle algorithm. Percentage of mineralized tissue was calculated with respect to the total explant volume. CTvox (Bruker micro-CT, BE) was used to create 3D visualization.

### DNA quantification, RNA isolation and gene expression analysis

DNA assay kit QuantiT dsDNA HS kit (Invitrogen) was used to quantify the DNA content from cell lysate according to the manufacturers protocol. RNA was isolated from the lysate with the RNeasy Mini Kit (Qiagen) according to the manufacturers protocol and quantified with NanoDrop 2000 (Thermo Scientific). RevertAid H Minus First Strand cDNA Synthesis Kit (Thermo Scientific) was used for reverse transcription. 1 μg oligo(dT18) was added to 11 μL RNA for 5 min at 65 °C, the reaction mixture (4 μL 5× reaction buffer, 1 μL ribolock ribonuclease inhibitor, 2 μL dNTPmix (10 × 10−3 m), and 1 μL RevertAid H Minus M-MuL VRT) was added, cDNA was generated using the Applied Biosystems Veriti 96-Well Fast Thermal Cycler (60 min at 42 °C followed by 10 min at 70 °C) and diluted in RNase-free water to 5 ng mL-1. Fast SYBR™ Green Master Mix (Thermo Scientific), 5 ng mL-1 cDNA and specifically designed primers were used to perform Quantitative reverse transcription polymerase chain reaction (qRT-PCR) consisting of a denaturation step at 95 °C, followed by 40 cycles of 95 °C, 3 s and 60 °C, 20 s. For quality control, a melting curve was generated between 60 °C and 99 °C. Gene expression data is presented relative to the housekeeping gene hypoxanthine-guanine phosphoribosyltransferase (GAPDH) and relative to day 0 monolayer culture.

### Histological stainings

Microtissues were gently flushed out from their microwells, concentrated, and fixed in 2 % PFA overnight, mixed in 3 % agarose, dehydrated and embedded in paraffin overnight. Ectopic explants were fixed in 4 % PFA for 4 hours, decalcified in ethylenediaminetetraacetic acid (EDTA)/PBS (pH 7.5) for 10 solution changes at 4 °C, dehydrated, and embedded in paraffin overnight, and sectioned at 5 μm thickness. Before histological staining, the slides were deparaffinized in Histoclear (Laborimpex).

For Safranin O/Fast green (Sigma) staining, sections were deparaffinized and dehydrated, counterstained with Hematoxylin (Merck, cat 6525) for 1 minute, briefly dipped in acid alcohol (1% HCL in 70 % EtOH), rinsed in water, stained with 0.03% Fast Green (KLINIPATH, cat 80051) and then dipped in 1% glacial acetic acid followed by 7-minute staining in 0.25 % SafraninO (KLINIPATH, cat 640780). Then the samples were washed in tap water, dehydrated with an ethanol series, cleared in histoclear, and mounted in Pertex for microscopy imaging. For Alcian Blue staining, sections were deparaffinized, rehydrated and stained with filtered Alcian Blue solution for 30 min at room temperature. After washing, the slides were counterstained by Nuclear Fast Red for 5 min before washing, dehydrating and mounting.

## Ethical statement

All procedures on animal experiments were approved by the local ethical committee for Animal Research, KU Leuven. The animals were housed according to the regulations of the Animalium Leuven (KU Leuven).

## Statistical analysis

All statistical analyses were performed using standard function in R (R core team). Statistical significance was defined at p < 0.05. Pairwise comparisons were done through a two-sided, unpaired t-test. Factor analysis was done by analysis of variance (ANOVA) for the model: defects = aspiration x dispension + time. Data is presented as mean and standard deviation from 4 samples. Symbols used are *p<0.05, **p<0.01, ***p<0.001, ****p<0.0001.

## Software and code availability statement

All software is defined in a docker container which is freely available at https://hub.docker.com/r/gnasello/spheroid-env and all notebooks used to generate the image processing workflow are available in Github (https://github.com/isaakdecoene/spheroid_classifier)

## Data availability statement

The data that support the findings of this study are available upon reasonable request from the authors.

## Author Contributions

ID contributed to the experimental design, analysis and manuscript preparation. GN contributed to the automatization of the image processing pipeline, the creation of a Docker environment and revision of the manuscript. RFMC, IVH and SRV contributed to the execution of experimental activities. AP contributed to the design, execution, and analysis of the first DoE. GNH contributed to the conceptualisation and experimental design. CV, LG, and FPL contributed to the conceptualization and design of the study. IP contributed to the conceptualization and design of the study, and revision of the manuscript.

## Acknowledgments

We thank Kathleen Bosmans for performing the in vivo mice experiments. This work was done in the context and with the support of members of Prometheus, the KU Leuven R&D division for skeletal tissue engineering (http://www.kuleuven.be/prometheus).

## Figures and tables

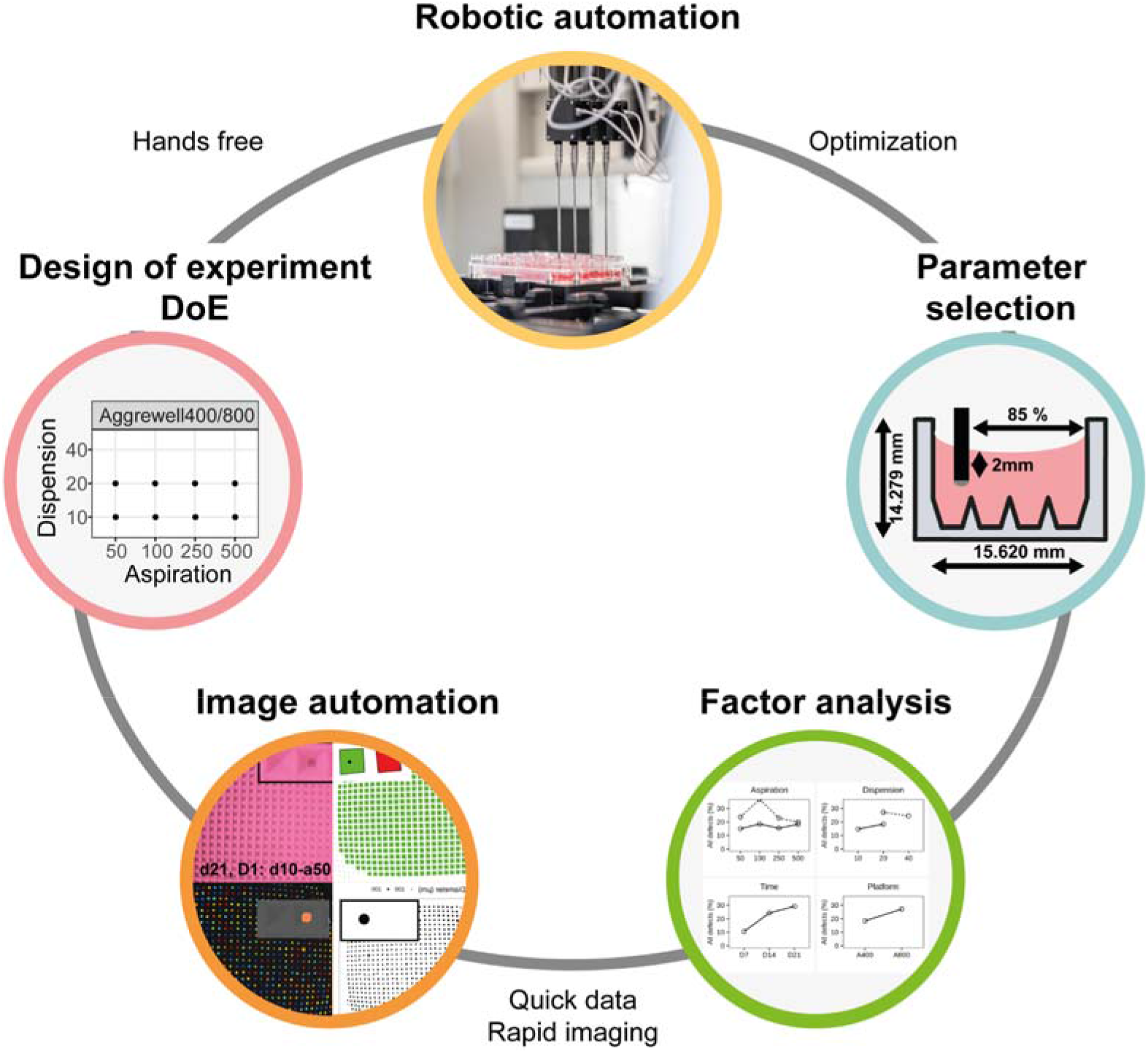

**Supplemental figure 1.**
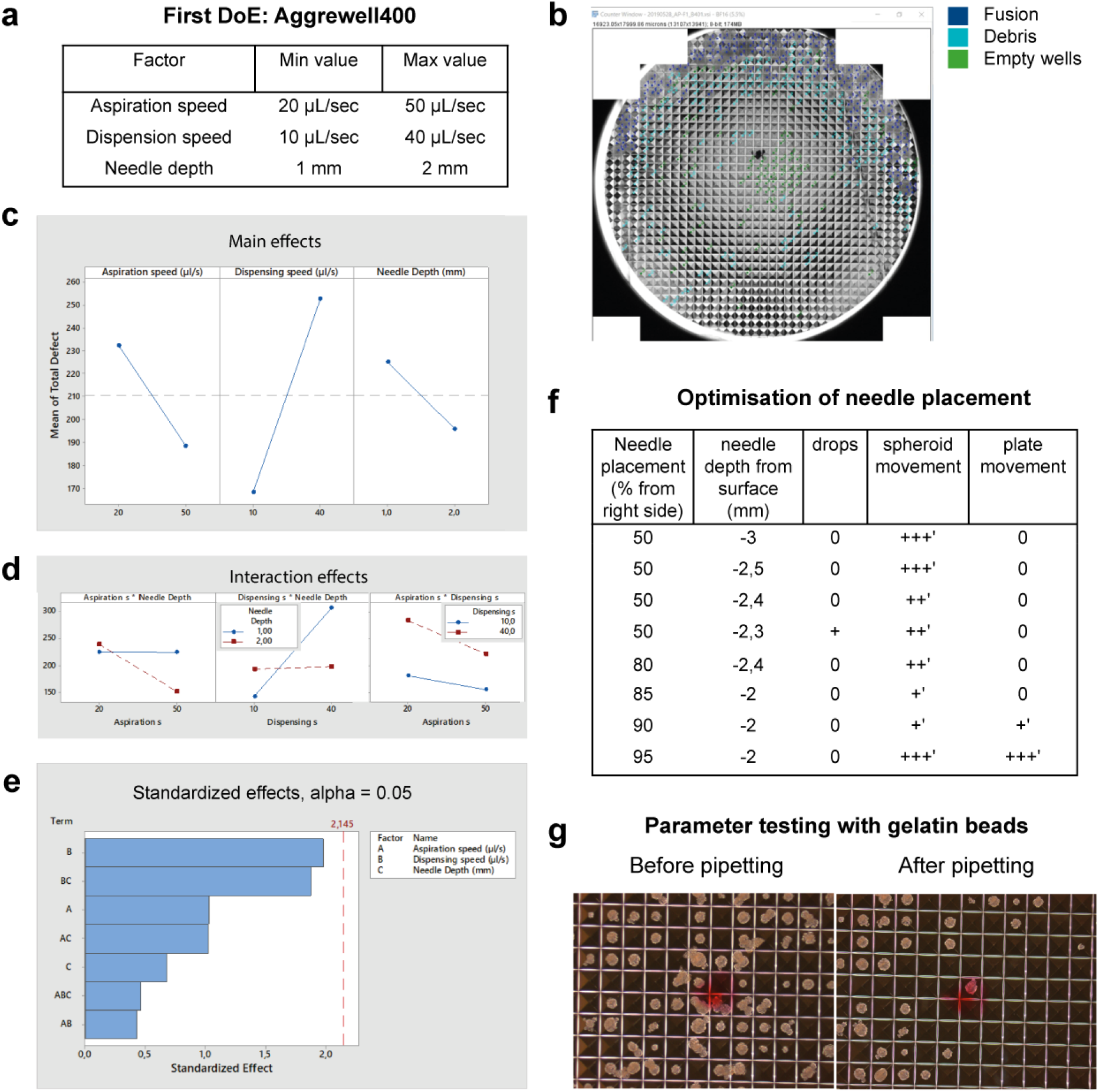
First parameter screening and optimisation. (a) DoE design showing factors and levels. (b) Example of manual analysis strategy. (c-e) DoE analysis results showing (c) main effects, (d) interaction effects, and (e) statistical significance. (f) Summary table of empirical testing of needle placement parameters. (g) Brightfield images of gelatine beads used during optimisation of needle placement with microwell size is 400 μm.

**Supplemental figure 2.**
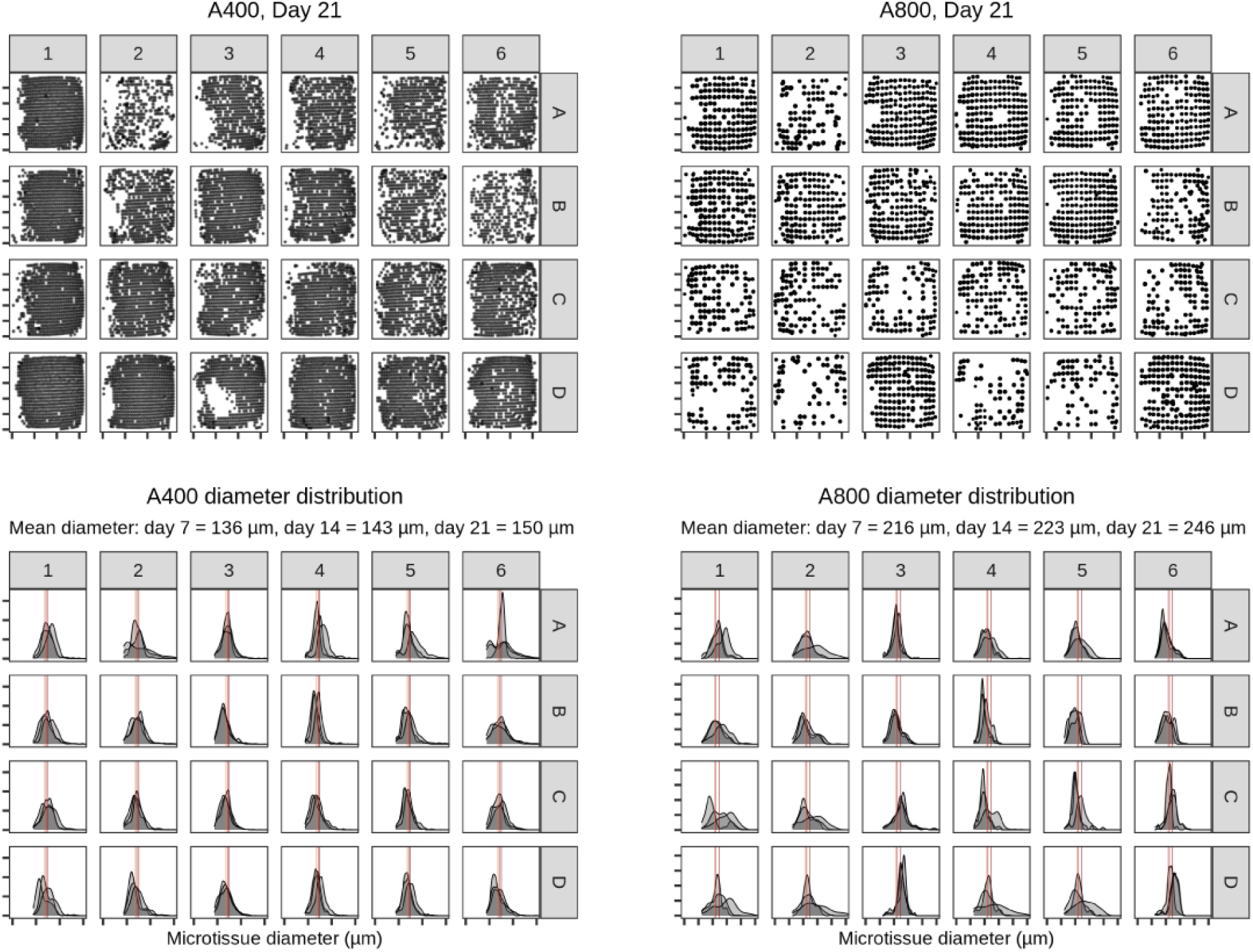
Microtissues as datapoints for process monitoring. (a-b) Microtissue locations and diameter (μm). (c-d) Density plots showing the diameter of microtissues in each well in the 24-well pate over time.

